# Subcontinental Genetic Diversity in the *All of Us* Research Program: Implications for Biomedical Research

**DOI:** 10.1101/2025.01.09.632250

**Authors:** Mateus H. Gouveia, Karlijn A. C. Meeks, Victor Borda, Thiago P. Leal, Fernanda S. G. Kehdy, Reagan Mogire, Ayo P. Doumatey, Eduardo Tarazona-Santos, Rick A. Kittles, Ignacio F. Mata, Timothy D. O’Connor, Adebowale A. Adeyemo, Daniel Shriner, Charles N. Rotimi

## Abstract

The *All of Us* Research Program (*All of Us*) seeks to accelerate biomedical research and address the underrepresentation of minorities by recruiting over one million ethnically diverse participants across the United States. A key question is how self-identification with discrete, predefined race and ethnicity categories compares to genetic diversity at continental and subcontinental levels. To contextualize the genetic diversity in *All of Us*, we analyzed ∼2 million common variants from 230,016 unrelated whole genomes using classical population genetics methods, alongside reference panels such as the 1000 Genomes Project, Human Genome Diversity Project, and Simons Genome Diversity Project. Our analysis reveals that participants within self-identified race and ethnicity groups exhibit a gradient of genetic diversity rather than discrete clusters. The distributions of continental and subcontinental ancestries show considerable variation within race and ethnicity, both nationally and across states, reflecting the historical impacts of U.S. colonization, the transatlantic slave trade, and recent migrations. *All of Us* samples filled most gaps along the top five principal components of genetic diversity in current global reference panels. Notably, “Hispanic or Latino” participants spanned much of the three-way (African, Native American, and European) admixture spectrum. Ancestry was significantly associated with body mass index (BMI) and height, even after adjusting for socio-environmental covariates. In particular, West-Central and East African ancestries showed opposite associations with BMI. This study emphasizes the importance of assessing subcontinental ancestries, as the continental approach is insufficient to control for confounding in genetic association studies.

## INTRODUCTION

Underrepresentation of racial and ethnic minorities in the United States (U.S.) has significantly limited biomedical studies, reflecting a broader global issue of underrepresentation of non-European ancestry populations. This lack of diversity hampers progress in developing precision medicine that effectively serves the nation’s highly diverse population^1^. Additionally, many genomic and biomedical studies in the U.S. are constrained by small sample sizes, lack of comprehensive phenotype data, and limited funding which impedes the ability to support long-term longitudinal research efforts. The *All of Us* Research Program (*All of Us*) aims to accelerate biomedical research and address minority underrepresentation by collecting comprehensive longitudinal health data on at least one million samples reflecting diversity across the U.S.^2^.

Recently, *All of Us* investigators published an analysis of the current release (version 7) of short-read whole-genome sequencing (WGS) data from 245,388 participants^3^. This study and others^3,4^ reported that Uniform Manifold Approximation and Projection (UMAP)^5^ recapitulated known patterns of population structure and that participants may cluster within discrete race and ethnicity categories, sparking debate on the validity and interpretability of these results^6,7^. UMAP^8^ is a nonlinear technique to reduce the complexity of multidimensional data and facilitate visualization in low-dimensional plots. This method is commonly applied in single-cell analysis and is designed to preserve the local structure of the data at the expense of the global structure. Consequently, differences can be exaggerated, and patterns that do not exist in the original data can be introduced. Since admixture generates populations with allele frequencies that are linear combinations of those in the parental populations, admixture is fundamentally a linear additive process.

There has been extensive discussion and recent recommendations regarding using race, ethnicity, and ancestry in scientific research^9,10^. Ancestry is defined as the population origin of an individual’s alleles at polymorphic sites^11^, which can be summarized as an average across the genome. In this study, we consistently adopt the genome-wide definition of ancestry. In contrast, race and ethnicity are socio-cultural or geopolitical constructs, with race associated with abstract beliefs about shared genetic ancestry or biological traits, and ethnicity connected to abstract perceived cultural practices^12^. In the questionnaire used by *All of Us*, racial categories primarily reflect politically recognized definitions of race within the country used by the U.S. Census, while ethnicity refers solely to whether a person is Hispanic or Latin American^13^. The definitions and use of race and ethnicity vary significantly across countries^14^, with examples like the different classifications in Brazil^15^, where race is often considered fluid, meaning it is a subjective perception of physical appearance and individuals may identify differently depending on social context, in contrast to the more rigid racial categories commonly used in the U.S. Similar variations can be observed across other countries, such as in Colombia and Mexico, where mestizo (mixed) identity is a dominant classification, despite the complex interplay of genetic and cultural diversity^15^. In the Netherlands, the country of birth indicator is widely accepted as a criterion for identifying ethnic groups^16^, while the concept of race is not utilized. Another complexity is that ethnic groups can be racialized, as exemplified by Hispanics/Latinos in the U.S., while racial groups can be ethnicized, as seen with Asians through shared cultural characteristics^10^. Despite the correlation between self-identified race/ethnicity and genetic ancestries, including in the U.S.^17,18^ and Latin America^19^, race and ethnicity are poor proxies for genetic ancestry^9^.

Based on classical population genetics methods, we assess the genetic diversity, admixture, and ancestry of 230,016 unrelated *All of Us* participants in the context of a newly compiled reference panel of genetic diversity. Our analyses address four key questions: (1) What does the population structure in *All of Us* look like across U.S. Census race and ethnicity population descriptors? (2) Does the genetic diversity in *All of Us* mirror the diversity seen in global diversity panels? (3) Is there variability in continental and subcontinental ancestries within racial and ethnic groups, and does this variability differ across U.S. geographical regions and states? (4) Are genetic ancestries associated with biological traits after accounting for major socio-environmental factors?

## RESULTS

### Genetic Diversity in All of Us

To assess the genetic diversity in *All of Us*, we analyzed the WGS data of 230,016 unrelated participants. Our preliminary analysis using all genetic variants in the WGS dataset proved computationally prohibitive, consistent with previous *All of Us* population genetics analyses^3^. Therefore, our analyses were based on a subset of variants from the WGS dataset; however, this subset included ∼2 million high-quality genome-wide variants across the 22 autosomes compared with ∼200,000 variants across two chromosomes used in a previous study^3^. Our analyses of genetic diversity rely on classical methods in population genetics, including principal component analysis (PCA)^20^, ADMIXTURE^21^, and *F* statistics^22^. We acknowledge that the genetic variation, population structure, and sample composition of the dataset influence unsupervised clustering analyses (e.g., PCA and ADMIXTURE).

Unsupervised PCA of *All of Us* participants identified five statistically significant (*p* < 0.05) principal components (PCs) with a genome-wide contribution of SNPs to the variance explained by each PC (SNP loading) and large eigenvalues (Figs. 1 and 2 and Fig. S1). These five PCs were used to interpret population structure within *All of Us*. PCs beyond PC 5 primarily reflected genetic differentiation due to variation in specific loci (*e.g.*, the *MHC* locus in PC 6 and both lactase [*LCT*] and MHC in PC 9), rather than broad genome-wide differentiation (Fig. S1). Participants were mapped with gradients of genetic diversity along PCs 1 and 2 (Fig. 1A and 1B), consistent with findings from biobank-level genomic studies of ancestrally diverse U.S. samples in New York City^23^. Interestingly, being born in the U.S. or elsewhere did not influence the overall pattern of population structure (Fig. S2). This suggests that recent migrants reflect the overall genetic diversity of *All of Us* participants born in the U.S. “Black or African American” participants represent a wide range of diversity along PC1 (Fig. 1A), while “Hispanic or Latino” participants are distributed across PCs 1 and 2 (Fig. 1B), encompassing nearly the whole triangle of space expected in three-way admixture.

**Fig. 1.**
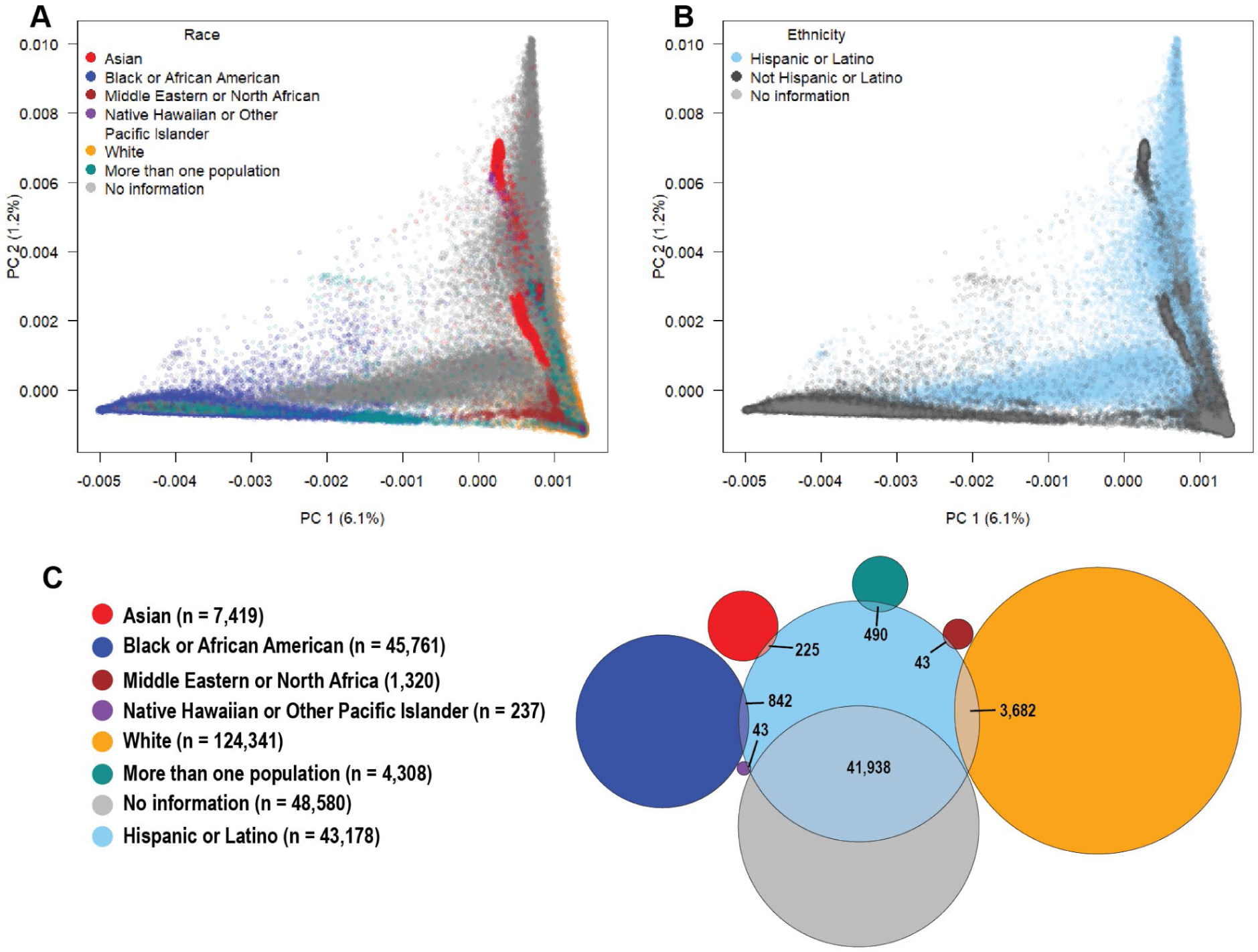
Genetic diversity in *All of Us* and the overlap between self-identified U.S. race and ethnicity. **A, B)** The first two principal components (PCs 1 and 2) of *All of Us* participants with race (**A**) and ethnicity (**B**) categories. **C)** Venn diagram showing the overlap between *All of Us* participants who self-identified with one race and as “Hispanic or Latino”.

**Fig. 2.**
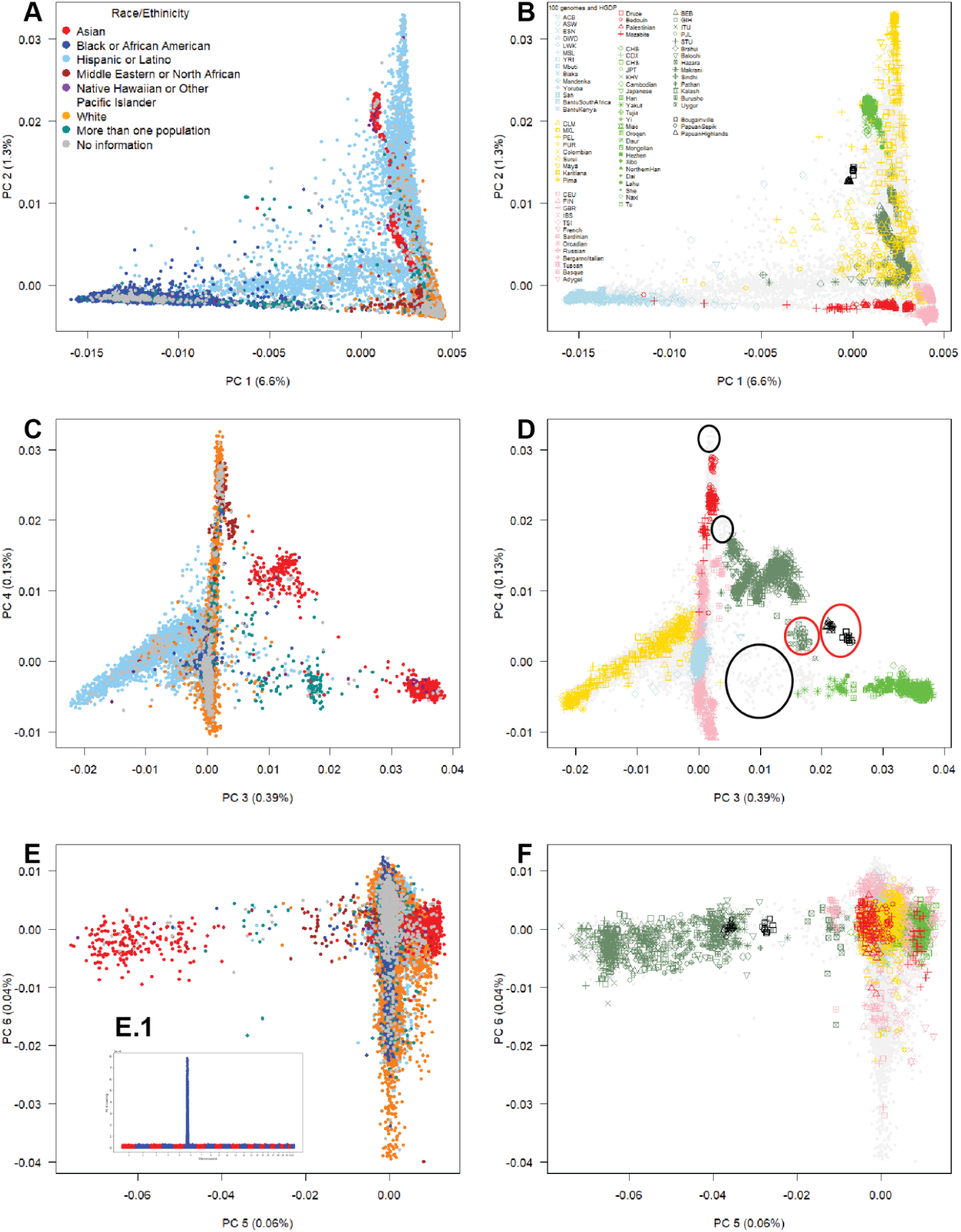
The breadth of genetic diversity in *All of Us* in the context of global genetic diversity. **A, B)** The first two principal components (PCs 1 and 2) in *All of Us* participants alongside our reference genetic diversity panel. **C, D)** The third and fourth components (PCs 3 and 4). **E, F)** The fifth and sixth components (PCs 5 and 6). The inset highlights that SNP loadings on PC 6 reflect genetic differentiation driven by the *MHC* locus, rather than genome-wide differentiation. **1kGP** = 1000 Genomes Project, **HGDP** = Human Genome Diversity Project. Race/ethnicity categories are integrated into the same PCA representation for visualization. The black circles represent genetic diversity in *All of Us* that is not captured by the reference panels, while the red circles represent genetic diversity within the reference panels that is absent in *All of Us*.

Most (92%) individuals who did not self-identify within any U.S. race category (Fig. 1A) self-identified ethnically as “Hispanic or Latino” (Fig. 1B), suggesting that the existing U.S. racial categories do not reflect the identity of this group. We observed varying degrees of cross-classification by race among those participants who self-identified ethnically as “Hispanic or Latino”: “White” (3,682), “Black or African American” (842), “Asian” (225), “Native Hawaiian or Other Pacific Islander” (43), and “More than one race” (490). This result reflects the complexity of how race and ethnicity categories are perceived in the U.S. by “Hispanic or Latino” individuals. The observation that “Hispanic or Latino” ethnicity includes individuals from all predefined categories of race reinforces how relying on these categories may provide insufficient adjustment for population structure in association studies.

Based on classical measures of genetic differentiation (*F*-statistics or fixation indices) between participants within race and ethnicity categories, we observed that most of the genetic variance is within race and ethnicity groups (1-*F_IT_* = 93.4%) rather than between groups (average *F_ST_* = 0.042). This result is consistent with studies of human genetic diversity^24–26^ showing that within-population differences account for most (84-95%) of the genetic variation, while differences among major geographic populations account for up to 5%. The average genetic differentiation between race and ethnicity groups (*F_ST_* = 0.042) is about half the average (*F_ST_*∼ 0.1) among populations across different continents^27,28^, suggesting that on average individuals assigned to worldwide populations will be more genetically distinct than individuals assigned to race or ethnicity.

### Multicontinental Genetic Diversity Represented by *All of Us*

To better explore the relationship between self-identification and gradients of genetic variation, we merged race/ethnicity categories into the same PCA representation (Fig. 2). We identified genetic diversity within *All of Us* by projecting samples from widely used reference panels (the 1000 Genomes Project^29^ and the Human Genome Diversity Project [HGDP]^30^, combined referred to as the “multicontinental diversity panel”) onto *All of Us* samples (Fig. 2). We observed that the genetic diversity in *All of Us* not only recapitulated but also exceeded the spectrum of diversity reflected in our multicontinental diversity panel across PCs 1 to 5 (Fig. 2).

“Black or African American” participants were distributed along PC 1, representing a gradient between African and European genetic diversity. “Hispanic or Latino” participants were spread across the top two PCs, reflecting the same gradient of African and European genetic diversity along PC 1 in addition to a gradient of European and Native American genetic diversity along PC 2. Also, we observed clustering of “Hispanic and Latino” participants distributed along a gradient of Native American genetic diversity defined by PCs 3 and 4. Importantly, *All of Us* samples project to spaces not covered by the 1000 Genomes Project or HGDP, including a broader range of three-way admixture patterns not observed in the Admixed American or Native American reference samples (PCs 1 and 2). Conversely, some genetic diversity within HGDP is not represented in *All of Us* (PCs 3 and 4), such as Central Asians (Hazara and Uygur) and Oceanians (Bougainville, Papuan Sepik, and Papuan Highlands).

### *All of Us* Coverage of Multicontinental Genetic Diversity

To evaluate the coverage of genetic diversity in *All of Us* in the context of multi-continental genetic diversity, we created a comprehensive panel capturing the genetic diversity, including 162 populations across all continents from different population genomics studies referred as to “global diversity panel”: the harmonized 1000 Genomes Project and the Human Genome Diversity Project (HGDP)^31^, and the Simons Diversity Project (SGDP)^32^, which provides a much broader globally sampling of indigenous populations^32^. PCA analysis of our global diversity panel showed different gradients of genetic diversity (Fig. 3): African *vs.* European (PC 1), European *vs.* Native American (PC 2), Asian *vs.* Native American (PC 3), Eastern *vs.* Southern Asian (PC 4), South African hunter-gatherer *vs.* other African populations (PC 5), and Northern European *vs.* Southern European and Middle Eastern (PC 6).

**Fig. 3.**
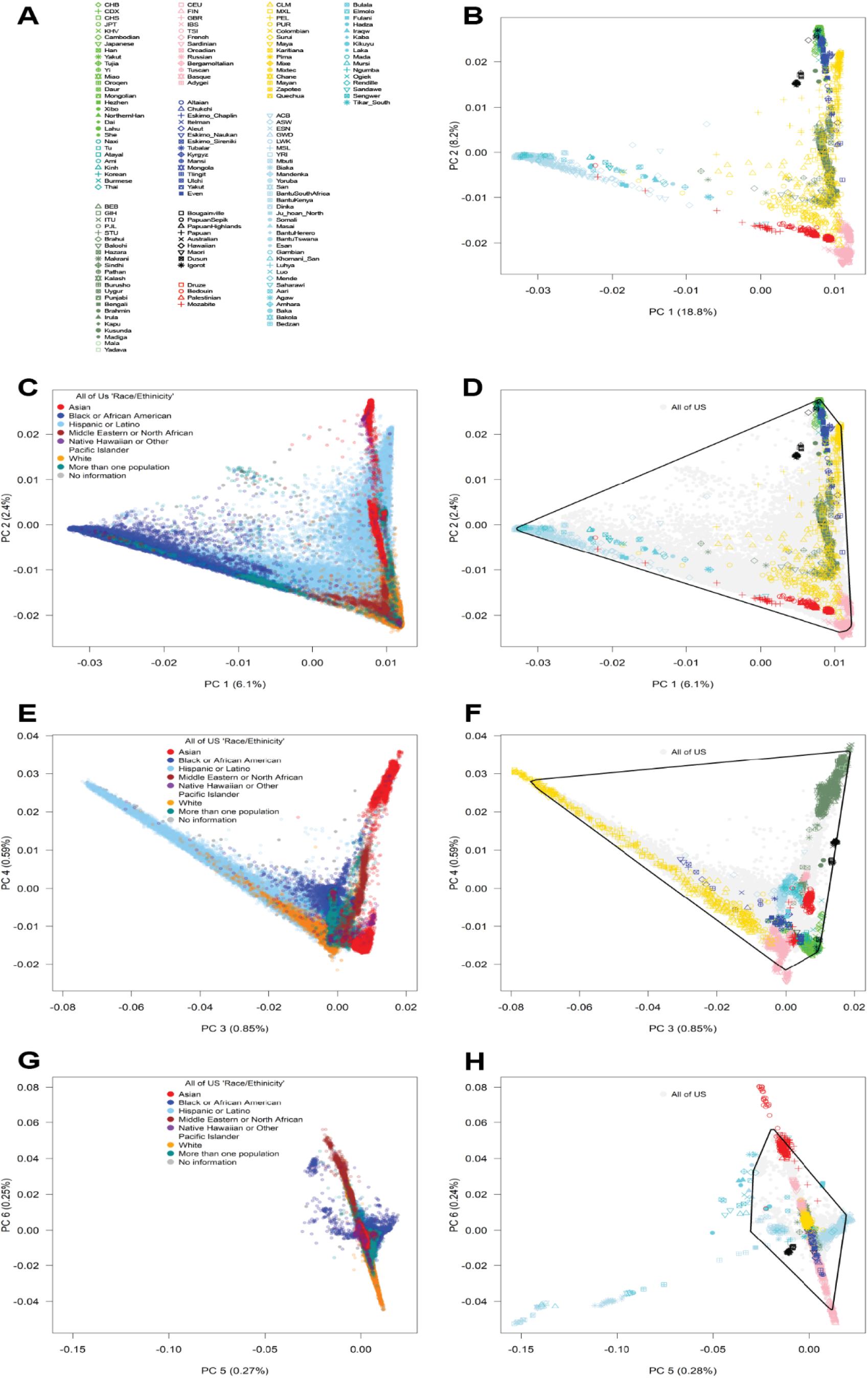
Coverage of multicontinental genetic diversity by *All of Us*. **A)** Population labels and **B)** the first two principal components (PCs 1 and 2) in our panel of multicontinental genetic diversity. **C, D)** The first two principal components (PCs 1 and 2) in *All of Us* participants alongside our reference genetic diversity panel. **E, F)** The third and fourth components (PCs 3 and 4). **G, H)** The fifth and sixth components (PCs 5 and 6). Convex hull areas in the PC plots represent genetic diversity coverage, with race/ethnicity categories integrated into the same plot. Panels C, E, and G highlight *All of Us* data whereas B, D, F, and H highlight our global diversity panel.

Projection of the *All of Us* samples onto our global diversity panel, along with the calculation of convex hull PC areas covered by *All of Us*, revealed substantial coverage of multi-continental diversity across five PCs (PCs 1–4 and 6; Fig. 3). Similar to our findings based on unsupervised PCA (Figs. 1 and 2), *All of Us* diversity not only covered most of PCs 1 and 2 but also extended to fill nearly the entire triangular space of three-way admixture. For PCs 3 and 4, *All of Us* samples covered broad genetic diversity, including a wider set of admixture combinations than typical two-way or three-way ancestries. Despite this extensive coverage, *All of Us* did not overlap with some populations, such as specific Native American populations (*e.g.*, the Karitiana in Brazil), specific European populations (*e.g.*, Sardinians and Basques, PC 3 *vs* PC 4), Middle Eastern populations (*e.g.*, Bedouin, PC 6), and Southern Asian populations (*e.g.*, Indian Telugu in the UK, PC 6). We confirmed that “White” participants covered most North-to-South European genetic diversity (∼90%) of our panel of European subcontinental diversity (Fig. S3)^33^. Consistent with previous reports on the genetics of the African diaspora^34,35^, the genetic diversity in *All of Us* did not capture the diversity specific to African hunter-gatherer populations (PC 5). Our analysis suggests that *All of Us* captures much of the worldwide genetic diversity in humans, while covering a broader range of admixture than is represented in current reference panels.

### Ancestry and Admixture in *All of Us*

To assess admixture in *All of Us*, we first performed unsupervised ADMIXTURE^21^ analysis using our global diversity panel. This analysis identified the most likely number of ancestral clusters as 13 (Fig. S4 and Table S1). Next, we projected *All of Us* samples onto our global diversity panel (supervised ADMIXTURE analysis) to estimate individual ancestry proportions across these 13 ancestry clusters (Fig. 4A and B). Dendrogram analysis of genetic differentiation, based on estimates of *F_ST_*among the ancestry clusters, revealed six clades of ancestries that we contextualize in terms of present-day geographic distributions (Fig. 4C): 1) West Central, West, South, and East African, 2) North European, South European, and Middle Eastern, 3) Indian and South Asian, 4) North Asian and Southeast Asian, 5) Native American, and 6) Oceanian. These clades of ancestries are consistent with estimated ancestry clusters reflecting major geographic regions observed in classical studies of human genetic diversity^26,24^.

**Fig. 4.**
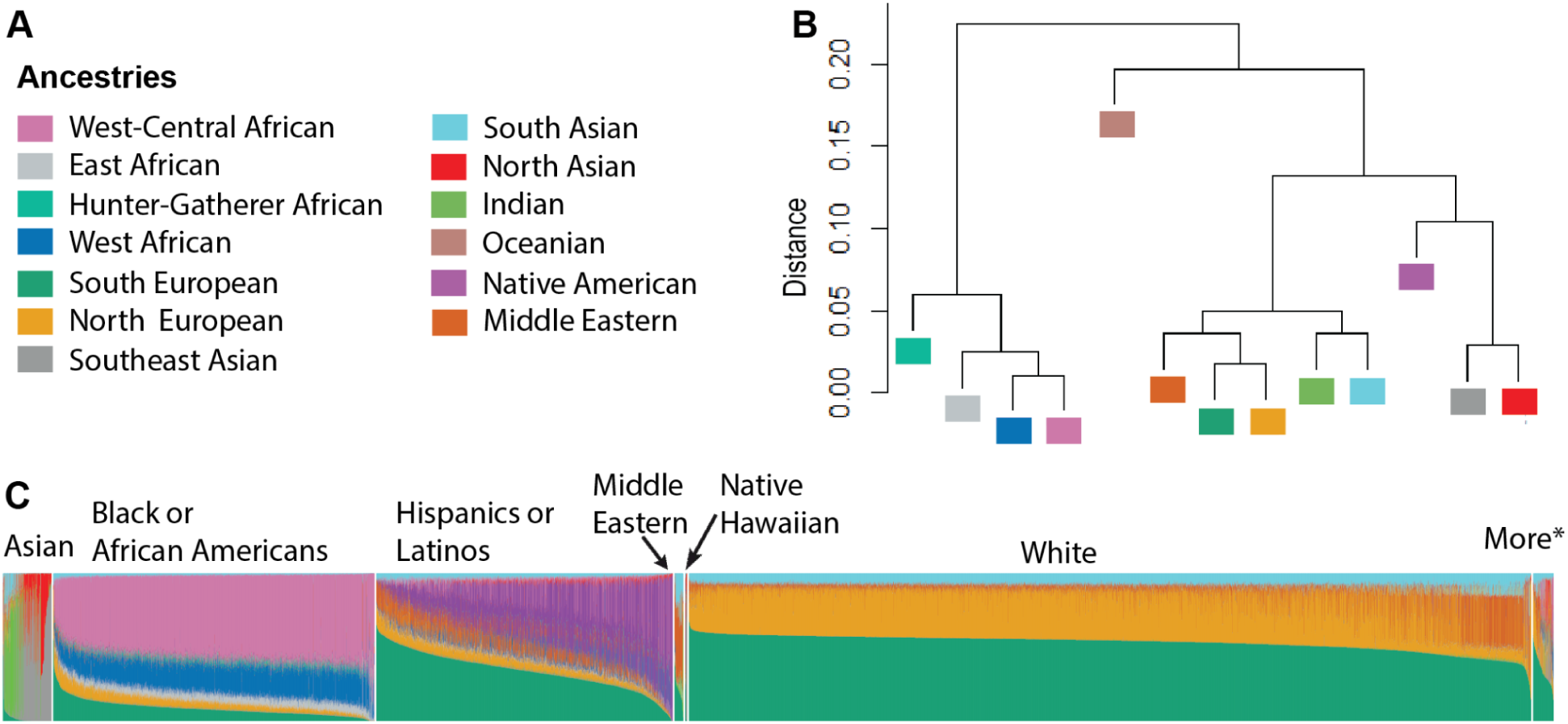
Continental and subcontinental ancestries in *All of Us*. **A)** Color legend representing the 13 most likely ancestry clusters identified from a global genetic diversity reference panel. **B)** Dendrogram of genetic relationships among the 13 ancestry clusters, based on estimates of *F_ST_*. **C)** Bar plot displaying individual ancestry proportions in *All of Us* participants with self-identified race and ethnicity categories, inferred from ADMIXTURE projection analysis. *More = *All of Us* participants self-identified in more than one race.

We observed large variability in individual ancestry proportions within race and ethnicity categories (Figs. 4 and 5, Tables S3 and S4). “Hispanic or Latino” participants had mean ancestral proportions consistent with three-way admixture (Table S2): total European (49.95%, SD = 19.15), Native American (31.35%, SD = 22.59), and African (13.47%, SD = 17.30, Table S2). Most “Hispanic or Latino” participants (75.1%) had at least 50% total African, European, or Native American (Table S3). “Black or African American” participants had mean ancestral proportions mostly consistent with two-way admixture (Table S2): 82.72% (SD = 10.87) African and 14.31% (SD = 9.35) European. However, we observed “Black or African American” participants with ≥ 50% total European (n = 483, 1.06%) ancestry as well as other ancestries (Table S3).

**Fig. 5.**
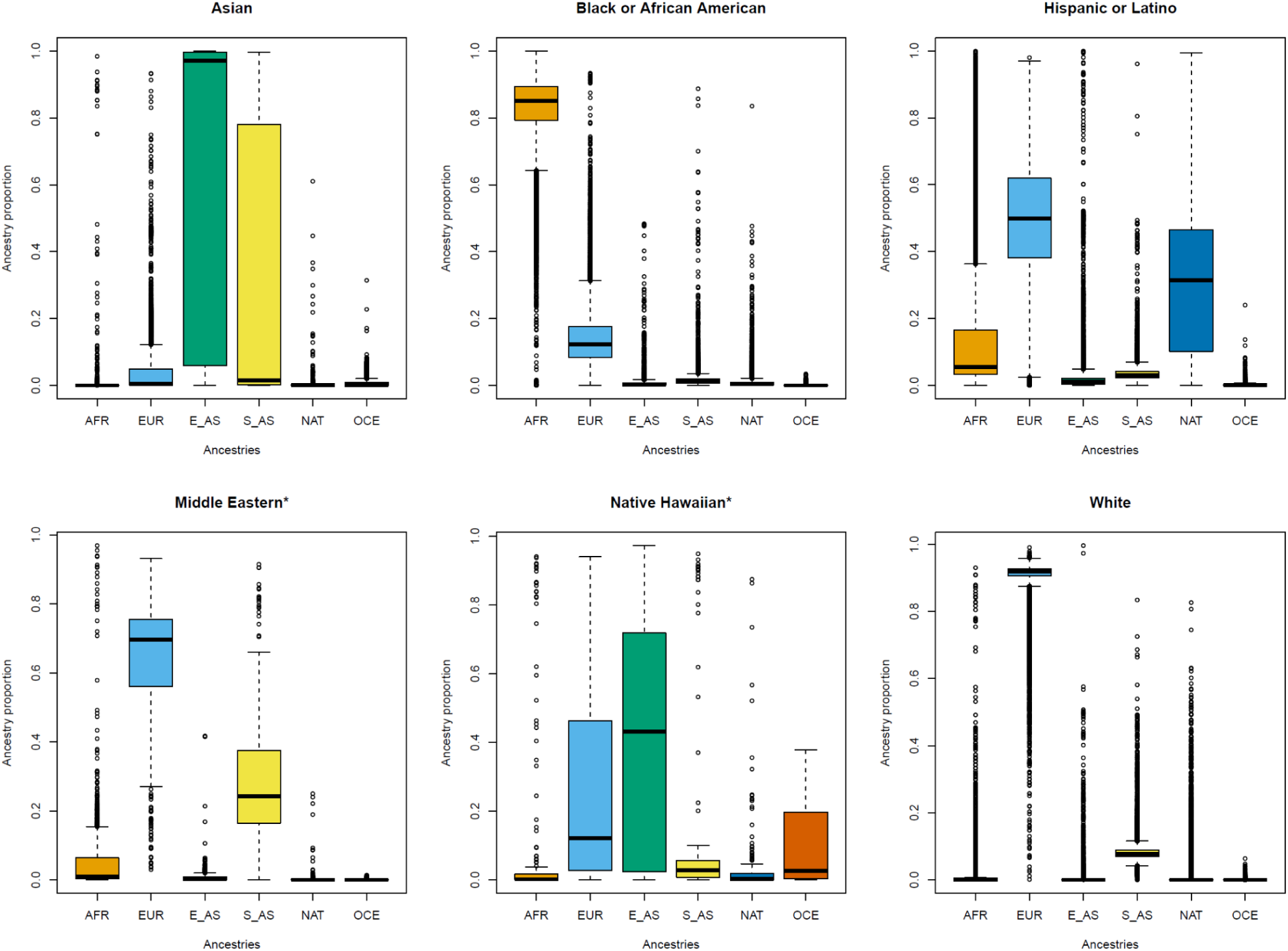
Distribution of continental ancestry proportions in *All of Us* participants within race and ethnicity categories. A) Asian, **B)** Black or African American, **C)** Hispanic or Latino, **D)** *Middle Eastern = Middle Eastern or North African, **E)** *Native Hawaiian = Native Hawaiian or Pacific Islander, and **F)** White. AFR = African, EUR = European, E_AS = East Asian, S_AS = South Asian, NAT = Native American, OCE = Oceanian.

“White” participants primarily had European ancestries (90%, SD = 5.58), with a minor proportion of South Asian ancestry (8.38%, SD = 2.85, Table S2). South Asian ancestry likely reflects admixture from ancient migrations of southern steppe peoples^36^. Additionally, some “White” participants were found to have ≥ 50% African (n = 29, 0.23%), Native American (n = 103, 0.08%), and other ancestries (Table S3). “Middle Eastern or North African” participants were primarily mixtures of European (65.59%, SD = 14.62) and South Asian (26.49%, SD = 15.30) ancestries (Table S2). “Asian” participants predominantly had East Asian (68.35%, SD = 42.76) and South Asian (24.73%, SD = 38.00) ancestries. “Native Hawaiian or Other Pacific Islander” participants exhibited a highly admixed ancestry profile, with a notable proportion of East Asian ancestry (37.14%, SD = 32.74, Table S2). We also observed a wide distribution of subcontinental ancestries within race and ethnicity groups. For example, while the majority of ‘White’ participants (n = 120,673; 97%) had predominantly Southern European ancestry, a subset (n = 3,418; 3%) exhibited predominantly Northern European ancestry, which is strongly associated with Finnish and Estonian European populations.

### Regional Variability in the *All of Us*

When assessing the distribution of ancestries across U.S. states, we observed regional variation. Among “Black or African American” participants (Fig. 6), mean total African ancestries was higher in southern states, such as South Carolina (85.6%) and Florida (85.3%) compared to western states like California (77.3%) and Arizona (79.0%). African subcontinental ancestries largely reflected the overall pattern of total African ancestries, with a notable exception in South Carolina, where West African ancestry was slightly elevated. East African ancestry, however, was more uniformly distributed across U.S. states. This alignment between total African ancestries and subcontinental patterns is likely influenced by the historical transatlantic slave trade, which predominantly involved individuals from West African regions^37^.

**Fig. 6.**
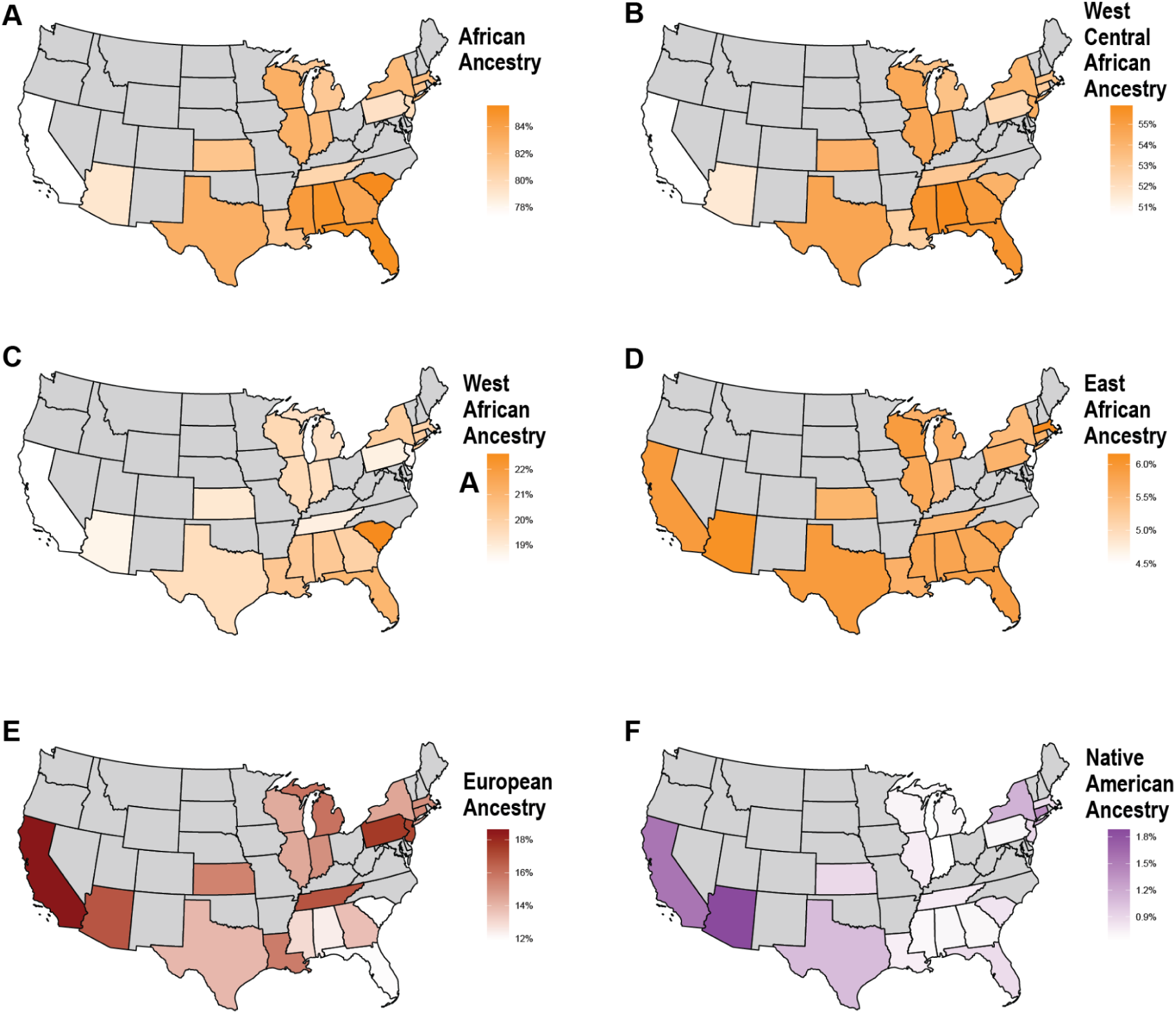
The distributions of continental and subcontinental ancestries in self-identified “Black or African American” participants by state. **A)** Total African, **B)** West-Central African**, C)** West African, **D)** East African, **E)** European, and **F)** Native American ancestry proportions. States with less than 50 individuals are excluded in gray.

For “Hispanic or Latino” participants (Fig. 7), ancestry patterns were highly diverse across U.S. states. The highest mean Native American ancestry proportions were observed in southwestern and western states, including California (45.6%), Texas (39.4%), and Arizona (38.0%), while certain southeastern states, such as South Carolina (50.1%) and Tennessee (45.0%), also showed elevated levels. In contrast, “Hispanic or Latino” participants had lower Native American ancestry proportions in northeastern states such as Pennsylvania (16.3%), New York (16.6%), and New Jersey (18.1%), but also in Florida (17.4%). “Hispanic or Latino” participants with the highest total European ancestry proportions were found in Florida (64.3%) and Pennsylvania (63.0%), followed by New Mexico (58.3%) and New Jersey (55.3%). Total African ancestry in “Hispanic or Latino” participants exhibited an increasing gradient from west to east, with the highest proportions observed in New York (31.5%) and the lowest in New Mexico (3.9%), California (5.4%), and Arizona (5.6%).

**Fig. 7.**
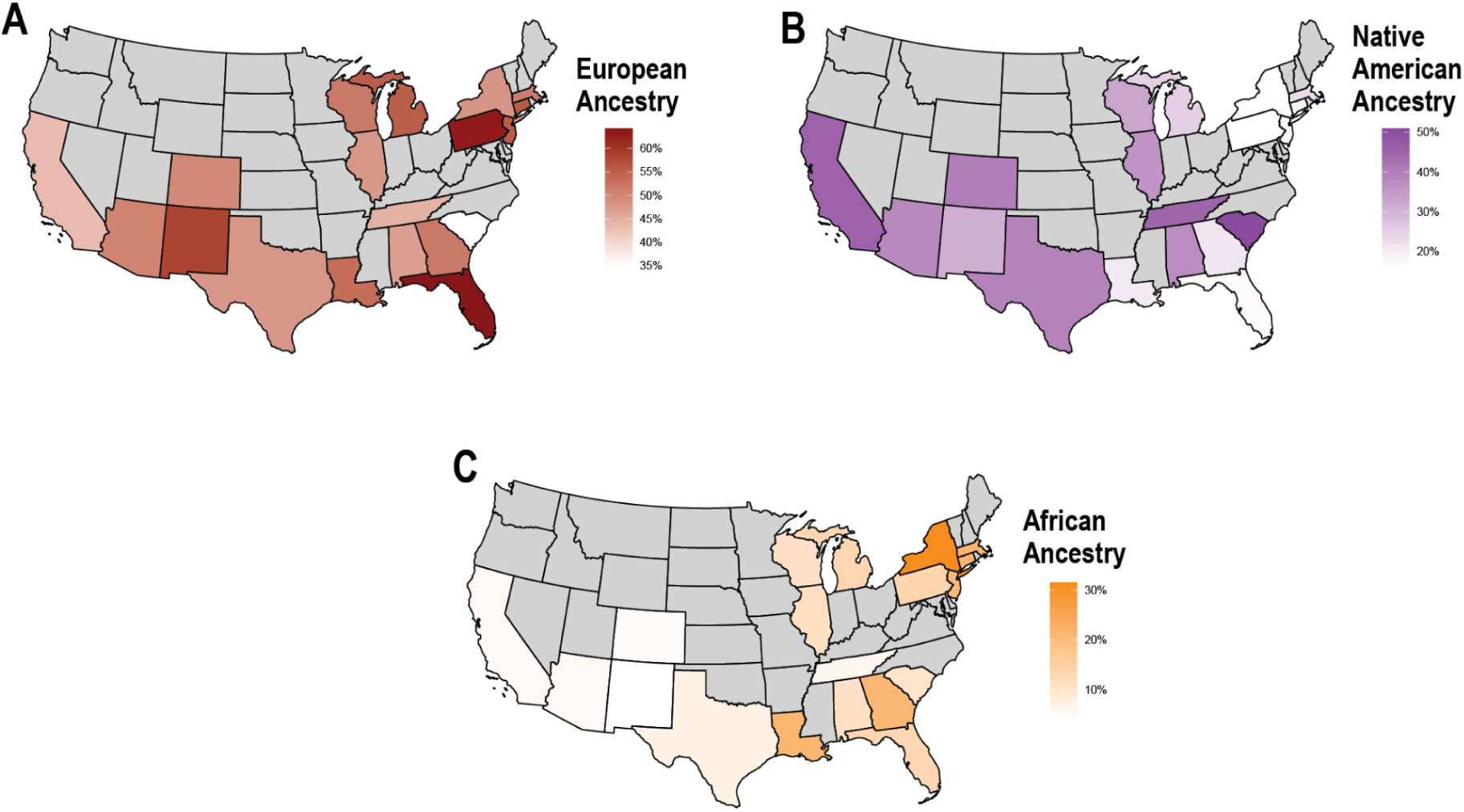
The distributions of continental ancestries in self-identified “Hispanic or Latino” participants by state. **A)** European, **B)** Native American, and **C)** African ancestry proportions. States with less than 50 individuals are excluded in gray.

The mean total European ancestry of “White” participants (Fig. 8) was higher in northeastern states, while lower proportions were found along the Mid-Atlantic and southern states. Subcontinental European ancestry also varied by state, with North European ancestry more prevalent in Midwestern states and South European ancestry more prevalent in the South. Notably, Middle Eastern ancestry showed a sharp contrast, with mean proportions significantly higher in New York and New Jersey (∼15%) compared to states like Alabama and Arkansas (∼3%).

**Fig. 8.**
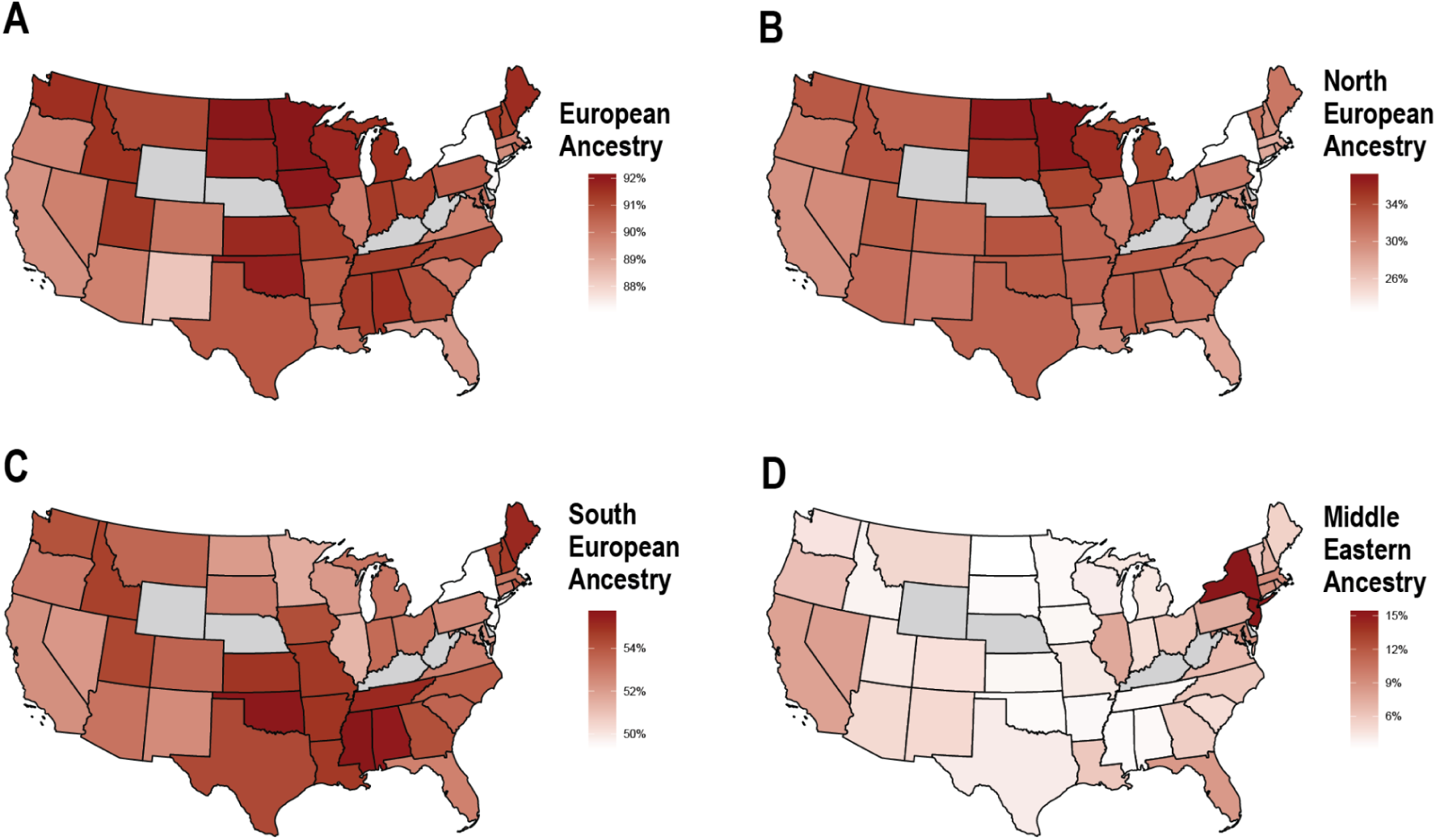
The distributions of European and Middle Eastern or North African continental and subcontinental ancestries in self-identified “White” participants by state. **A)** Total European, **B)** North European, **C)** South European, and **D)** Middle Eastern or North African ancestry proportions. States with less than 50 individuals are excluded in gray.

### *All of Us* Participants in Race/Ethnicity and Predicted Ancestry Groups

Genetic epidemiology studies in the U.S. often conduct association analyses, including genome-wide association studies (GWAS), by stratifying participants according to self-identified race and ethnicity^38–40^. However, in the *All of Us* Research Workbench, which includes GWAS data on approximately 3,400 phenotypes in the “All by All tables”, participants are stratified by categorical ancestry groups (based on ancestry estimation) rather than self-identified race and ethnicity. As GWAS findings in the All by All tables have been used to replicate previous associations from studies designed based on self-identified race/ethnicity, we examined the correspondence between participants stratified by race/ethnicity and genetic ancestry.

We compared the ancestry distributions of “Black or African American” and “White” participants with predicted African and European ancestry participants, respectively (Fig. 9A and B). We noticed that individuals stratified based on predicted categorical ancestry groups had more homogeneous ancestry distributions, with fewer participants with outlier profiles of ancestry compared with participants stratified by race (Fig. 9A and B). Despite similar ancestry distributions of participants stratified based on these approaches, we observed that participants stratified by African (Fig. 9C) and European (Fig. 9D) genetic ancestry are distributed across multiple race/ethnicity groups. For example, participants in the predicted African category include individuals self-identifying as “Hispanic or Latino” (n = 2,981) or as “more than one race” (n = 1,400). Similarly, participants in the predicted European category include individuals self-identifying as “Hispanic or Latino”, “More than one race” (n = 1,400), or “Middle Eastern or North African” (n = 510).

**Fig. 9.**
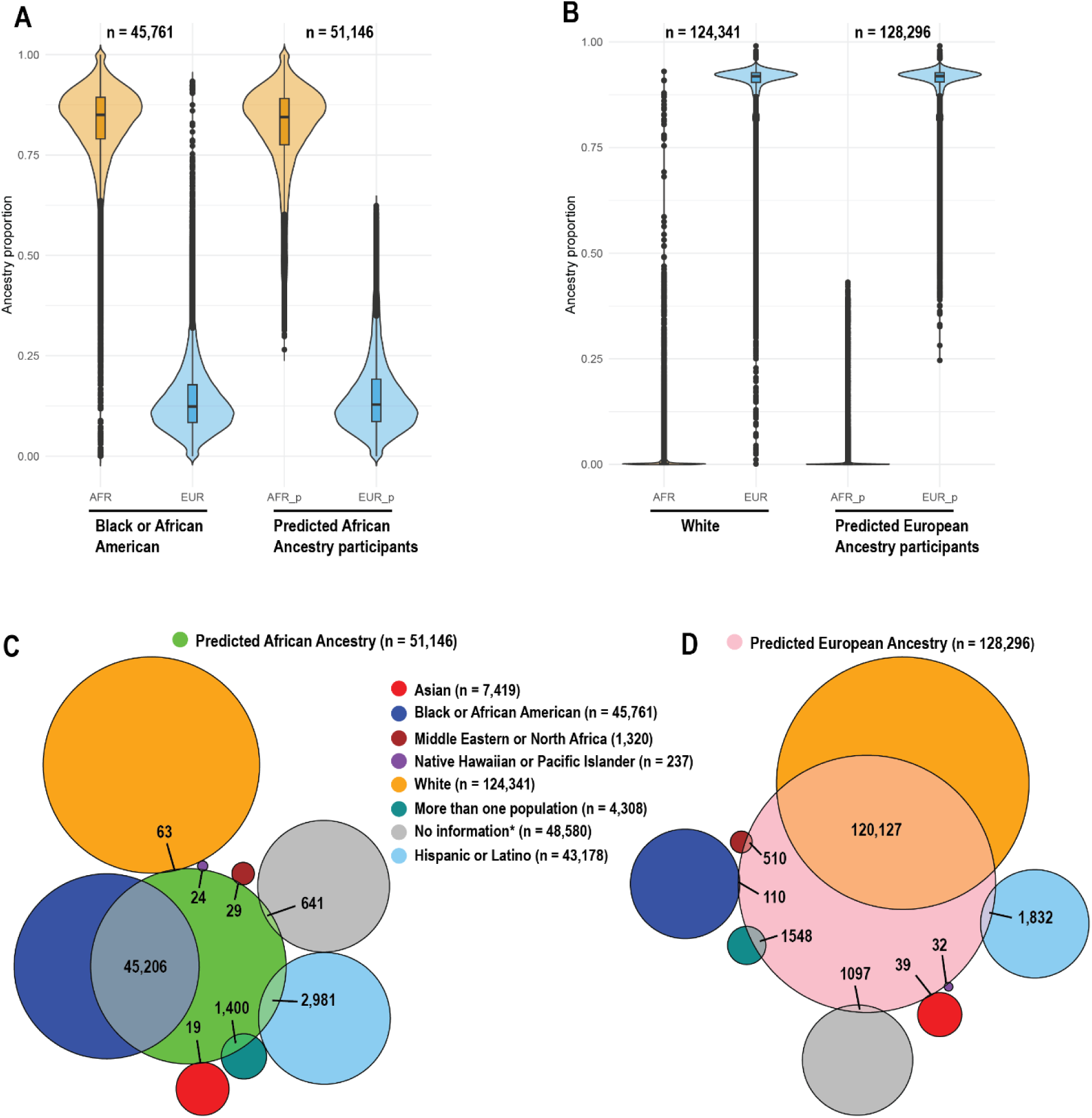
Distribution of *All of Us* participants by race/ethnicity and predicted genetic ancestry, highlighting overlaps between self-identified race/ethnicity and genetic ancestry predictions. **A)** Violin and box plots comparing African ancestry distribution among participants self-identifying as “Black or African American” with those stratified by predicted African genetic ancestry. **B)** Violin and box plots comparing European ancestry distribution among participants self-identifying as “White” with those stratified by predicted European genetic ancestry. **C)** Venn diagram illustrating the overlap between predicted African genetic ancestry and self-identified race/ethnicity categories. **D)** Venn diagram illustrating the overlap between predicted European genetic ancestry and self-identified race/ethnicity categories.

Overall, subsets of participants classified by race/ethnicity and ancestry prediction are not entirely concordant. Association studies, including GWAS and GWAS meta-analyses, should carefully consider these differences, especially since race/ethnicity may capture important socio-cultural or environmental effects.

### Genetic Ancestries and Association with Biological Traits in *All of Us*

To evaluate the impact of ancestry on biological traits, we assessed associations between ancestry clusters in *All of Us* individuals and body mass index (BMI) and height. Given the substantial variability in genetic ancestry (both nationally and at the state level) within race/ethnicity and the inadequacy of these categories as proxies for genetic ancestry, we did not stratify individuals by race/ethnicity in our association analyses. Instead, we included race/ethnicity alongside comprehensive social and environmental covariates to adjust the association models. To mitigate the effects of data sparsity, we performed the association analyses including individuals with at least 10% of the corresponding continental ancestry (or the combined subcontinental ancestries within that continent; Table S4 and S5).

First, we evaluated the association between genetic ancestry and height (Table S4), a trait that varies significantly across populations and has a large narrow-sense heritability (> 80%)^41^. In Europe, for example, height follows a pronounced south-to-north gradient, with taller statures more commonly observed in northern regions^41^. Consistent with previous studies^33,42^, North European ancestry was strongly associated with greater height (β = 0.107, SE = 0.004, *p*-value = 4.35 × 10^–204^) whereas both South European Ancestry and Middle Eastern ancestry were associated with lower height (β = -0.028, SE = 0.004, *p*-value = 2.21 × 10^–14^). West-Central African ancestry was associated with greater height (β = 0.045, SE = 0.004, *p*-value = 4.35 × 10^–29^) while Hunter-Gatherer African ancestry was associated with lower height (β = -0.196, SE = 0.043, *p*-value = 5.22 × 10^–6^). West and East African ancestries were not significantly associated with height. Both Indian, South Asian, and Southeast Asian ancestries were significantly associated with lower height whereas North Asian ancestry was associated with increasing height. In line with previous reports^43^, Native American ancestry was strongly associated with lower height (β = -0.088, SE = 0.002, *p*-value = 1.40 × 10^–274^).

Next, we evaluated the association between ancestry and BMI (Fig. 10 and Tables S5). We found that Asian and European ancestry clusters were significantly associated with lower BMI, whereas Native American ancestry was associated with higher larger BMI (β = 0.009, SE = 0.003, *p*-value = 1.56 × 10^–3^). Notably, West-Central African ancestry was associated with higher BMI (β = 0.019, SE = 0.005, *p*-value = 1.80 × 10^–05^) while East African ancestry was associated with lower BMI (β = -0.041, SE = 0.014, *p*-value = 3.18 × 10^–3^). Consistently, BMI polygenic scores (PRS), adjusted to account for differences due to ancestral background, revealed a positive correlation between West-Central African ancestry and PRS estimates for BMI (*r²* = 0.078, p = 0.0001). In contrast, East African ancestry showed a negative correlation with BMI PRS estimates (*r²* = -0.033, p = 0.10). These results indicate that subcontinental ancestries can have opposite effects on biological traits and diseases. BMI polygenic scores (PRS)

**Fig. 10.**
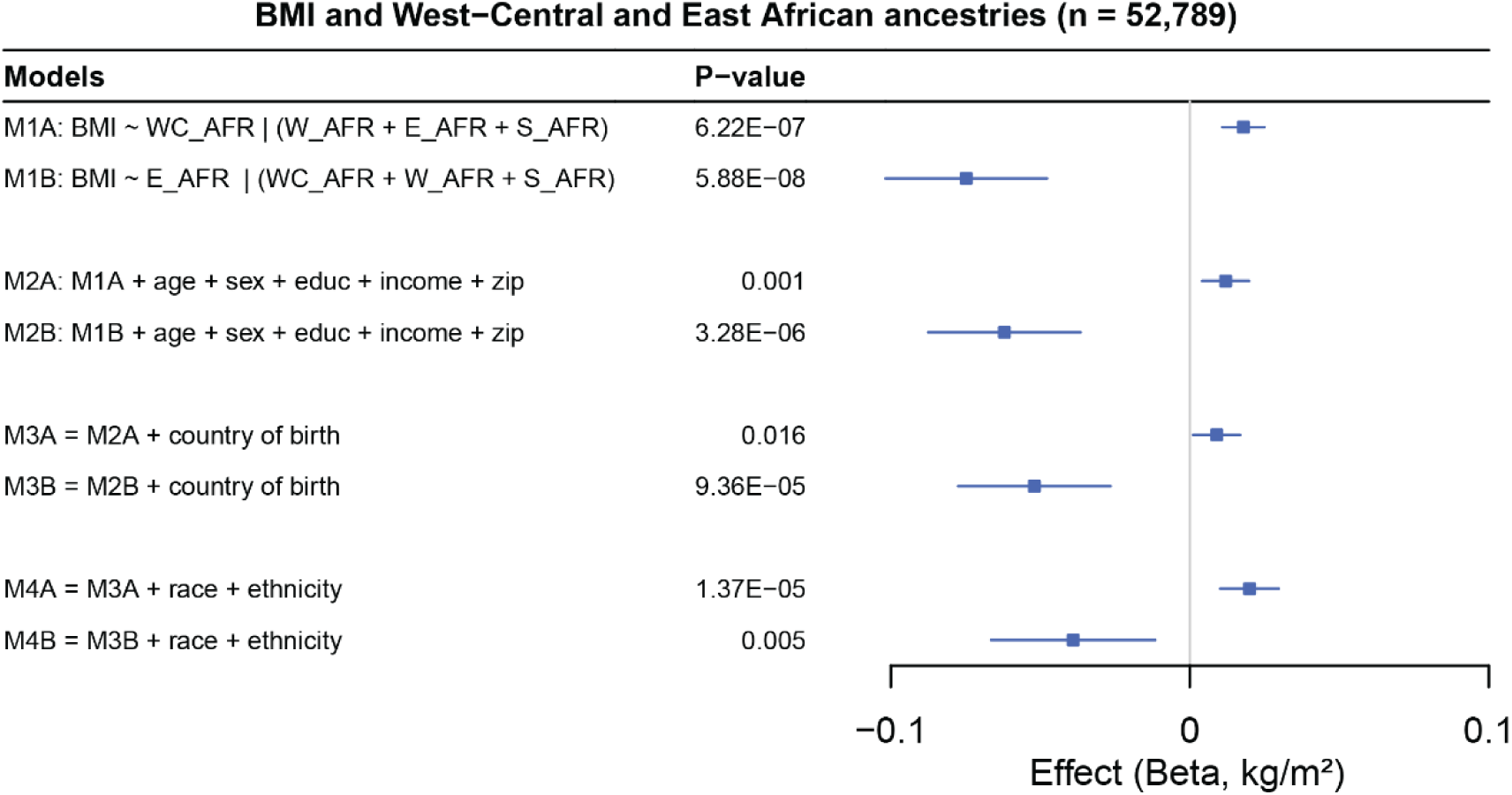
Forest plots showing the association between BMI and multiple traits, accounting for different levels of control of confounders across four different models. WC_AFR = West Central African, W_AFR = West African, E_AFR = East African, S_AFR = South African. *All of Us* participants with at least 10% of the combined subcontinental African ancestries were included.

Overall, associations between ancestry and traits were attenuated upon the inclusion of socio-cultural or environmental factors (Fig. 10 and Tables S5 and S7). Furthermore, after adjusting for a set of covariates including 3-digit zip codes, the inclusion of both race/ethnicity and country of birth systematically improved model fit. This result suggests that while race and ethnicity should not be used as proxies for genetic population structure, they may capture additional environmental effects not typically accounted for by standard covariates in association models. Although race/ethnicity may serve as a proxy for environmental effects, directly adjusting for more specific environmental factors affecting the outcome is preferable when such data are available^44^. Additionally, the country of birth may account for the effects of recent migration patterns to the U.S. and should be considered in association models using the *All of Us* dataset.

As a sensitivity analysis, we tested the associations between ancestry and BMI and height among participants with at least 50% of the corresponding continental ancestry (or combined subcontinental ancestries within that continent; Tables S6 and S7). Most results were consistent with the 10% threshold approach. However, including race/ethnicity as a covariate led to a poorer model fit for participants with at least 50% Native American ancestry, as the vast majority (99.6%) self-identified as ‘Hispanic or Latino’.

## DISCUSSION

By analyzing whole-genome sequencing data on 230,016 unrelated *All of Us* participants with a new panel of global genetic diversity, we conducted the largest population genomics analysis of U.S. samples that reflect the nation’s genetic diversity.

To assess genetic diversity in *All of Us* in the context of multicontinental populations, we created two panels of genetic diversity: the multicontinental diversity (1kGP, HGDP) and the global diversity (1kGP, HGDP, and SDGP). We employed two complementary approaches: first, an unsupervised PCA of the *All of Us* data, onto which the multicontinental diversity panel was projected; and second, an unsupervised PCA of the global diversity panel, onto which the *All of Us* data were projected. Our findings demonstrate that *All of Us* not only captures much of the diversity represented in existing reference panels but also bridges gaps along the first five axes of genetic diversity within these panels. This is consistent with the fact that ∼50 million people living in the U.S. are migrants from all continents and nearly every country in the world^45^. Our results align with findings from the Bio*Me* Biobank, a diverse multi-ethnic cohort from New York City (NYC)^23^, that identified that NYC individuals partially reflect the genetic diversity in populations from multiple countries around the world. This diversity offers important opportunities to improve the representation of populations previously excluded from genomics research.

Unlike previous analyses UMAP analyses, which suggested that participants in race/ethnicity categories may be distributed in discrete clusters^3,4^, we found gradients of genetic diversity that cut across those categories, consistent with findings from Bio*Me^23^*. Notably, *All of Us* “Hispanic or Latino” participants spanned nearly the entire spectrum of genetic diversity along the first two principal components. Most “Hispanic or Latino” (92%) did not self-identify within any specific race category, and those (8%) who did were represented across all possible pre-defined racial categories. Latin America has a recent history of admixture among multiple Native American, African, and European populations^17,34,46,47^. Furthermore, racial descriptors in Latin America are more fluid, whereas the U.S. has a more binary classification of “white” vs. “non-white”^15,48^.

Our results build upon findings from previous studies using smaller datasets^17,23,49,50^ that African, Native American, and European ancestries vary among individuals in the U.S. We further revealed significant variation in subcontinental ancestries, including pronounced regional differences across the country. These results highlight the complexity of genetic backgrounds and admixture within these groups and demonstrate that social constructs of race and ethnicity do not accurately reflect underlying genetic ancestry. Therefore, we do not recommend using race and ethnicity as proxies for ancestry in genetic studies, including association models. Rather, we support the use of race and ethnicity as markers of social, environmental, and historical factors that influence health outcomes^9^, when more direct measurements of these markers are not available^44^.

We observed that ancestral proportions within race/ethnicity categories vary by state. Most of the observed geographical variation in ancestries within racial and ethnic categories might be attributed to the history of colonization, the transatlantic slave trade, and recent migration patterns in the U.S. Among Black or African American participants, African subcontinental ancestries mirrored the total African ancestry, especially for West-Central African ancestry, which is the predominant African ancestry in the Americas^34^. This alignment between total African ancestry and subcontinental patterns is likely shaped by the U.S. slave trade, in which ancestry is traced to West-Central and West African countries such as Nigeria, Ghana, Benin, Ivory Coast, The Gambia, and Senegal^37^. This contrasts with the African ancestry profile in Brazil, where a more diverse array of African ancestries can be traced to regions including West, South, and East Africa (*e.g.*, Nigeria, Ghana, Angola, and Mozambique)^34^. An exception to this pattern was observed in South Carolina, where West African ancestry was elevated compared to other states. This observation is consistent with previous reports of a significant Grain Coast ancestry among African American men in South Carolina^51^, and aligns with the historical narrative of rice plantation farmers in South Carolina preferentially obtaining enslaved Africans from western Africa’s “Grain Coast” (*e.g.*, Senegal and Sierra Leone)^52^. “Hispanic or Latino” participants from U.S. Western states exhibited the highest proportion of Native American ancestry compared to other U.S. regions. This pattern aligns with the geographic proximity to Mexico, the historical context of western states (California, Texas, and Arizona) being part of Mexico until the mid-19th century, and recent migration patterns from Latin America. In contrast, “Hispanic or Latino” participants in New York had the highest proportion of African ancestry, consistent with the recent migration of Caribbeanns^53^, including Puerto Ricans to New York^54^. In agreement, a recent report highlighted differences in continental ancestries among Latin American immigrant groups in the U.S.^55^; for instance, Mexicans exhibited higher proportions of Native American ancestry, whereas Puerto Ricans showed greater levels of African ancestry. Among *All of Us* “White” participants (Fig. 6), the mean European ancestry was highest in the Northeastern U.S., while lower proportions were observed along the East Coast and in Southern states. This pattern is consistent with the settlement preferences of European migrants during the colonial period, influenced by favorable climates more similar to those of their European countries of origin^56^.

We demonstrated that classifying *All of Us* participants by race/ethnicity or genetic ancestry yields groupings that are not fully concordant. Consequently, using summary statistics from thousands of precomputed GWASs in the *All of Us* Researcher Workbench (“All by All tables”) for replication or meta-analyses in studies designed around race/ethnicity requires careful evaluation. One approach to address this issue is to adopt the same stratification strategy for *All of Us* participants as employed in the study being replicated or meta-analyzed.

We found that BMI and height are associated with genetic ancestries, even after adjusting models for key socio-cultural or environmental covariates such as age, sex, race, ethnicity, education, income, ZIP code, and country of birth. These results warrant new studies based on ancestry across the thousands of phenotypes available in *All of Us*, including admixture mapping^57^, a powerful gene mapping technique that leverages locus-specific ancestry to identify loci associated with differential risk or trait values by ancestry. Notably, our findings reveal that West-Central and East African ancestries are associated with BMI, but in opposite directions. Differential associations across African ancestries have also been shown for other health-related traits^58,59^. For instance, Meeks et al.^60^ demonstrated that triglycerides do not exert an equally strong association with other established risk factors in West and East Africans, underscoring the complexity of genetic and environmental factors influencing metabolic health. Our results and previous reports highlight the importance of avoiding using African continental ancestry as a single entity. Given this consideration and the well-documented subcontinental ancestries within Africa^34,61^, the Americas^34,55^, Asia^62^, and Europe^33^, along with their substantial regional variation within continents and in the U.S., we recommend treating continental ancestry not as a singular entity but as a composite of diverse subcontinental ancestries. Researchers should use plural forms, such as African, Native American, Asian, or European ancestries, and adjust association models to account for genetic diversity components that reflect subcontinental ancestries.

In conclusion, our analyses of the largest and most diverse U.S. biobank demonstrate that genetic diversity is characterized by distribution along a small number of gradients. Ancestry varies widely within race and ethnicity groups, both nationally and at the state level, emphasizing the inadequacy of such descriptors for defining genetically or biologically distinct populations. However, race and ethnicity may serve as proxies for capturing socio-cultural or environmental factors (*e.g.*, racism and social inequality) that are not typically accounted for by standard covariates in association models. The genetic diversity captured in *All of Us* reflects most of the diversity represented in public genetic reference panels, filling in gaps in existing panels. Furthermore, our findings highlight the importance of accurately characterizing subcontinental ancestries, as reliance on the continental ancestry approach is insufficient to control for confounding in association studies.

## METHODS

### Samples

We analyzed data from 230,016 unrelated *All of Us* participants with short-read whole-genome sequencing (WGS) available through the Curated Data Repository (CDR, Control Tier Dataset v7) in the *All of Us* Research Workbench. To establish a global multi-continental genetic diversity reference, we compiled genome-wide data from three multi-continental studies: the harmonized set of genomes^31^ from the 1000 Genomes Project (1kGP)^28^ and the Human Genome Diversity Project (HGDP)^30^ and genomes from the Simons Genome Diversity Project^32^ (Fig. 1A and Supplementary Data 1). The SGDP was designed to sample a broader range of globally diverse populations, with genomes sequenced from a few individuals (2-4) from previously underrepresented populations. We generated two reference datasets: The first comprising 3,433 unrelated individuals from 80 populations in the most used genetic diversity panels: 1kGP and the HGDP datasets (referred to as “multicontinental diversity panel”). The second, more comprehensive dataset combines 1kGP, HGDP, and SGDP, including 3,667 unrelated individuals representing 162 populations worldwide (referred to as “global diversity panel”).

### Data Curation

We used the Allele Count/Allele Frequency (ACAF) threshold files available in the *All of Us* Research Workbench. This call set includes approximately 99 million variants that are common in *All of Us* ancestry subpopulations, with population-specific allele frequencies (maf > 1%) or allele counts > 100. Our preliminary analysis revealed that current methods for assessing population structure^20,63–65^ using the entire *All of Us* set of variants and individuals were computationally inefficient or infeasible. Based on guidance from previous studies^21,34^, we thinned the dataset to approximately two million common SNPs. We performed data cleaning within and between datasets using PLINK 1.9^66^. Variants were filtered based on minor allele frequency (--maf 0.01), genotype missingness per variant (--geno 0.05), and genotype missingness per individual (--mind 0.05). Additionally, single nucleotide polymorphisms (SNPs) in linkage disequilibrium (LD) were pruned (--indep-pairwise 50 10 0.2).

### Relatedness

From 245,388 *All of Us* WGS participants, we generated a set of 230,016 unrelated samples by pruning a set of relatives (available in the *All of Us* Workbench^67^) optimized to maximize the set of unrelated samples. Briefly, relatedness was assessed using the pc_relate function implemented in Hail^68^ to estimate samples’ pairwise kinship coefficient (*Φ_ij_*). Pairs with *Φ_ij_* estimates above 0.1 were linked in a network framework to identify the largest set of unrelated individuals, minimizing the number of samples requiring pruning. Similarly, using our global multi-continental reference panel, we applied the KING method^69^ to estimate *Φ_ij_*. Genetic relationships were modeled as networks^70^, where individuals were linked if their *Φ_ij_*exceeded a threshold of 0.0884 (indicating first- and second-degree relatives^69^). Related individuals were then excluded using a maximum clique approach to minimize sample loss^70^.

### Population Structure and Genetic Ancestry

We performed supervised and unsupervised principal components analysis (PCA) using GCTA^64^ in the unrelated *All of Us* participants following two different approaches. First, to assess population structure in *All of Us* and identify known reference populations represented by *All of Us* diversity, we performed unsupervised PCA of *All of Us* followed by projection of our multi-continental diversity panel. We used ∼2 million SNPs not pruned by LD for this analysis. Second, to assess the coverage of *All of Us* participants of global genetic diversity, we performed unsupervised PCA using 535,933 independent variants in our global diversity panel followed by projection of the *All of Us* samples. We calculated the convex hull areas^71^ based on principal components to assess *All of Us* coverage of genetic diversity relative to our global diversity panel. For computational efficiency, unsupervised PCA on *All of Us* participants was performed using 10 random subsets of the entire dataset.

Genetic ancestry scores currently available in the *All of Us* workbench were determined using a trained classifier based on 16 principal components derived from gnomAD reference samples, based on 151,159 autosomal SNPs distributed on only two chromosomes. In our analysis to assess ancestry and admixture in *All of Us*, we used 535,933 independent variants distributed across the entire autosome. We performed unsupervised ADMIXTURE^21^ analysis using our global diversity panel to identify the most likely number of ancestral clusters in our panel. To evaluate the genetic relationship of the inferred ancestry clusters, we performed dendrogram clustering analysis of genetic relationships among the estimated ancestry clusters, based on estimates of *F_ST_*. Using the allele frequencies of the inferred ancestral clusters (ADMIXTURE projection analysis^21^), we projected *All of Us* participants onto our global diversity panel to estimate their ancestry proportions.

### Association Analysis

To assess the association of ancestry with BMI and height, we included all subcontinental ancestries within a continent (based on dendrogram *F_ST_* analysis) in the model simultaneously to evaluate the independent effect of each ancestry on the trait while controlling for the effect of the others. We performed the association analyses using four different models. Model 1 tested the association between the biological trait and subcontinental ancestries: *trait ∼ ancA_1_ + ancA_2_ + … + ancA_n_*, without adjusting for possible confounders. Model 2 included socio-economic covariates: *trait ∼ ancA_1_ + ancA_2_ + … + ancA_n_ + age + sex + education + income + zip code*. In model 3, considering that *All of Us* includes participants not born in the US, we added a covariate for recent immigration: *trait ∼ ancA_1_ + ancA_2_ + … + ancA_n_ + age + sex + education + income + zip code + country of birth*. In model 4, to account for socio-environmental effects captured by the U.S. Census categories, we added race and ethnicity: *trait ∼ ancA_1_ + ancA_2_ + … + ancA_n_ + age + sex + education + income + zip code + country of birth + race + ethnicity.* To mitigate the effects of data sparsity, we performed the association analyses including individuals with at least 10% of the corresponding continental ancestry (or the combined subcontinental ancestries within that continent). For sensitivity analysis, we repeated the associations analyses, but using participants with at least 50% of the corresponding continental ancestry (or combined subcontinental ancestries within that continent).

### Polygenic risk scores (PRS) estimates for BMI

Polygenic risk scores (PRS) for BMI were calculated for a subset of genetically diverse participants in the All of Us program (n = 13,475), who were previously selected to calibrate PRS estimates within the Electronic Medical Records and Genomics (eMERGE) network^72^. Briefly, the raw PRS was computed for all participants by summing the weighted risk alleles derived from previously published GWASs. Genetic ancestry was inferred based on the projection of participants into PC space. To mitigate biases introduced by variation in allele frequencies and linkage disequilibrium patterns across populations, a calibration model was developed using genetically diverse data from 13,475 *All of Us* participants. This model normalized the mean and variance of the PRS within ancestry groups, enabling the derivation of adjusted PRS values. Calibration ensured that PRS distributions were standardized, facilitating direct comparability across individuals irrespective of their continental genetic ancestry.

## ACKNOWLEDGMENTS

This work utilized the computational resources of the NIH HPC Biowulf cluster (https://hpc.nih.gov). The contents of this publication are solely the responsibility of the authors and do not necessarily represent the official views of the National Institutes of Health. The authors express their gratitude to the staff and participants of the All of Us Research Program. We thank Christopher Kachulis for insightful discussions on PRS estimates using eMERGE PRS pipeline.

## AUTHOR CONTRIBUTIONS

The project was conceived by M.H.G., D.S., C.N.R., and A.A.A. M.H.G. and D.S. assembled datasets. M.H.G. and D.S. analyzed genetic data. M.H.G., K.A.C.M., V.B., T.P.L., F.S.G.K., R.M., A.P.D., E.T.S., R.A.K., I.F.M., T.D.O’C., D.S., A.A.A., D.S., and C.N.R. contributed to data interpretation. M.H.G., K.A.C.M., A.A.A., D.S., and C.N.R. wrote the manuscript. All authors read the manuscripts and contributed with suggestions.

