## Supplemental Figures for "Subcontinental Genetic Diversity in the *All of Us* Research Program: Implications for Biomedical Research"

### Supplementary Figures

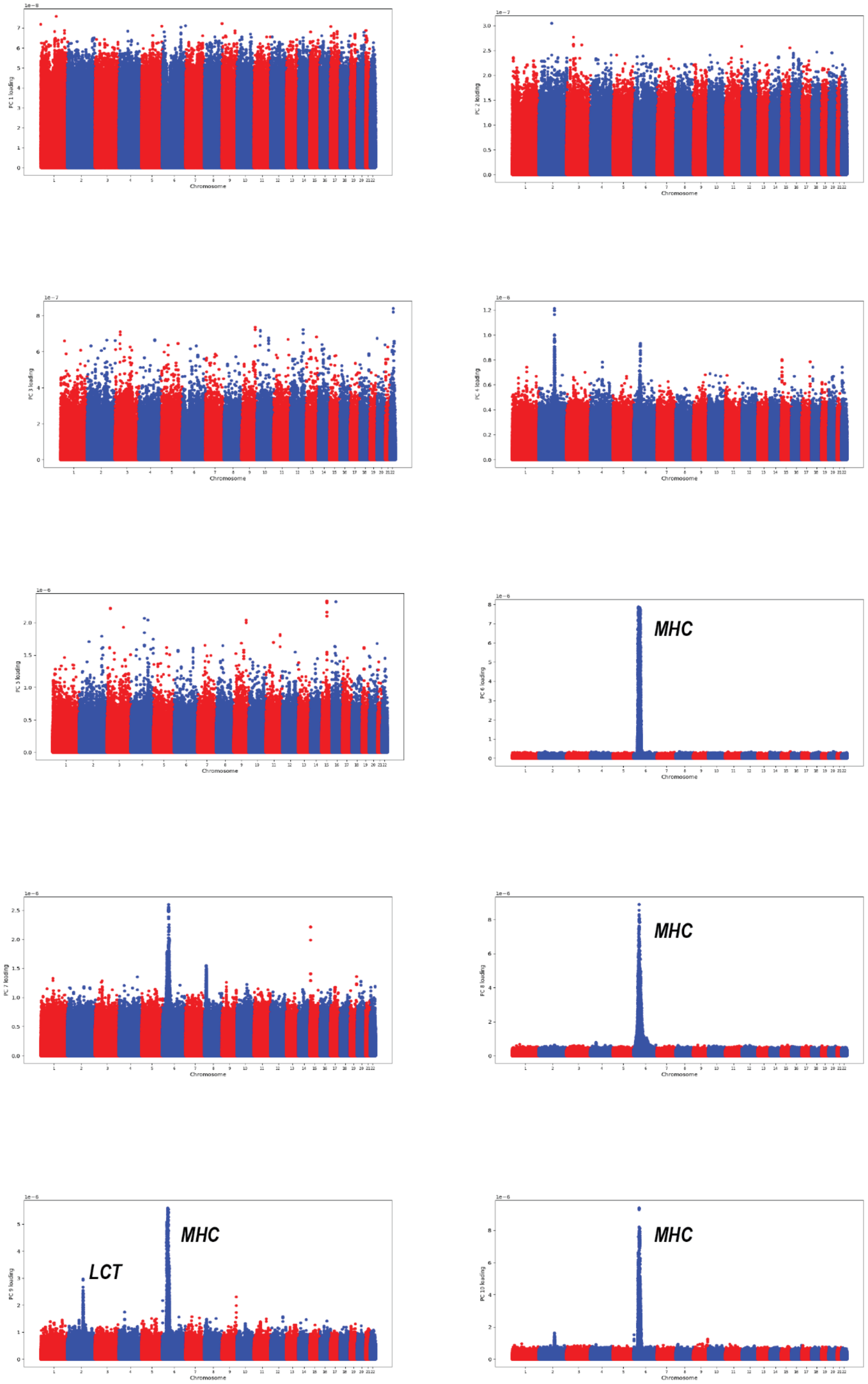


**Fig. S1. Manhattan plot of SNP loadings for the top 10 PCs from the unsupervised PCA analysis of *All of* Us.** *MHC,* Major Histocompatibility Complex. *LCT*, lactase.

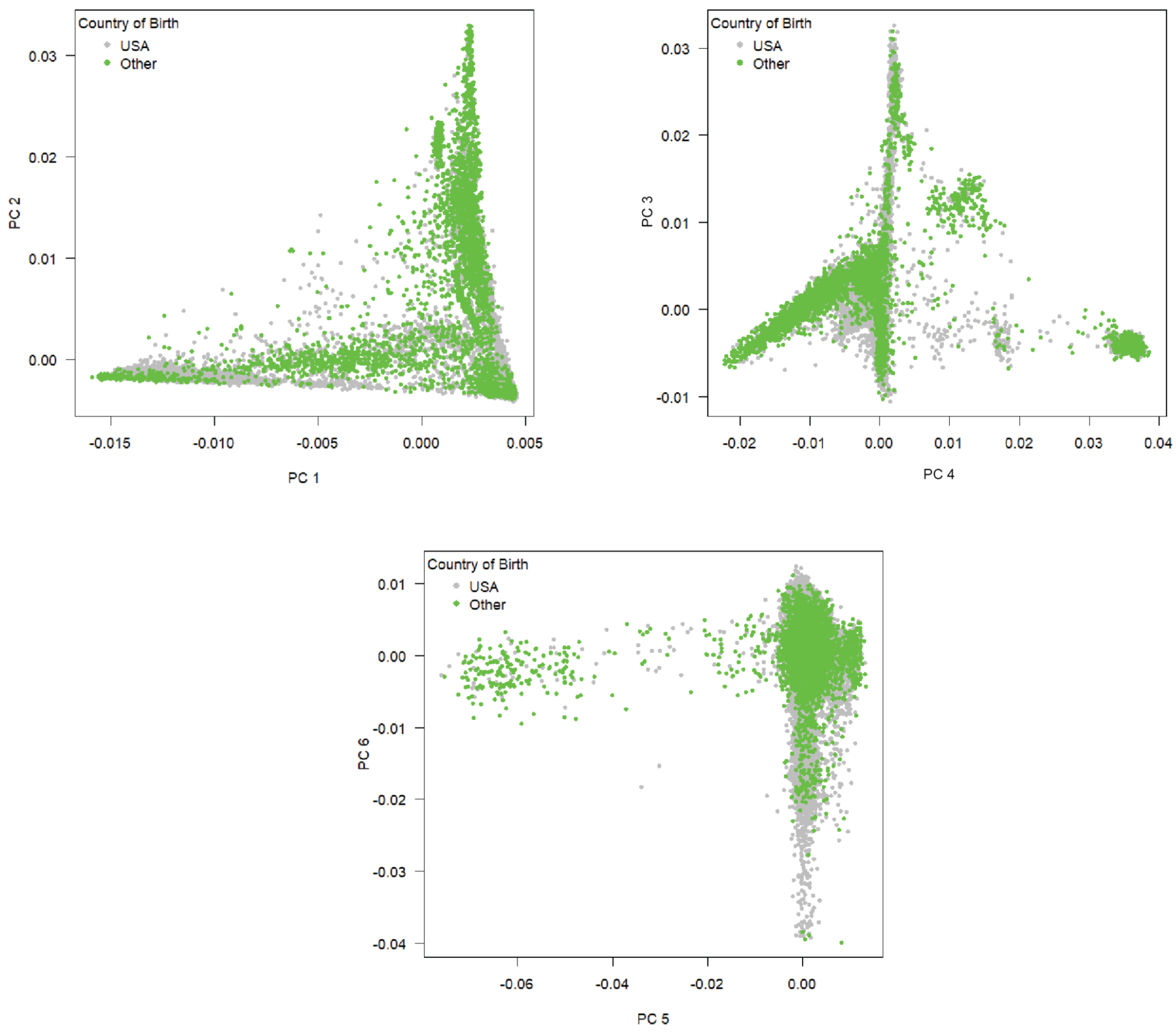


**Fig. S2. Genetic diversity of *All of Us* participants stratified by country of birth.** The *All of Us* workbench provides binary information on participants' country of birth, categorized as either "United States" or "non-United States".

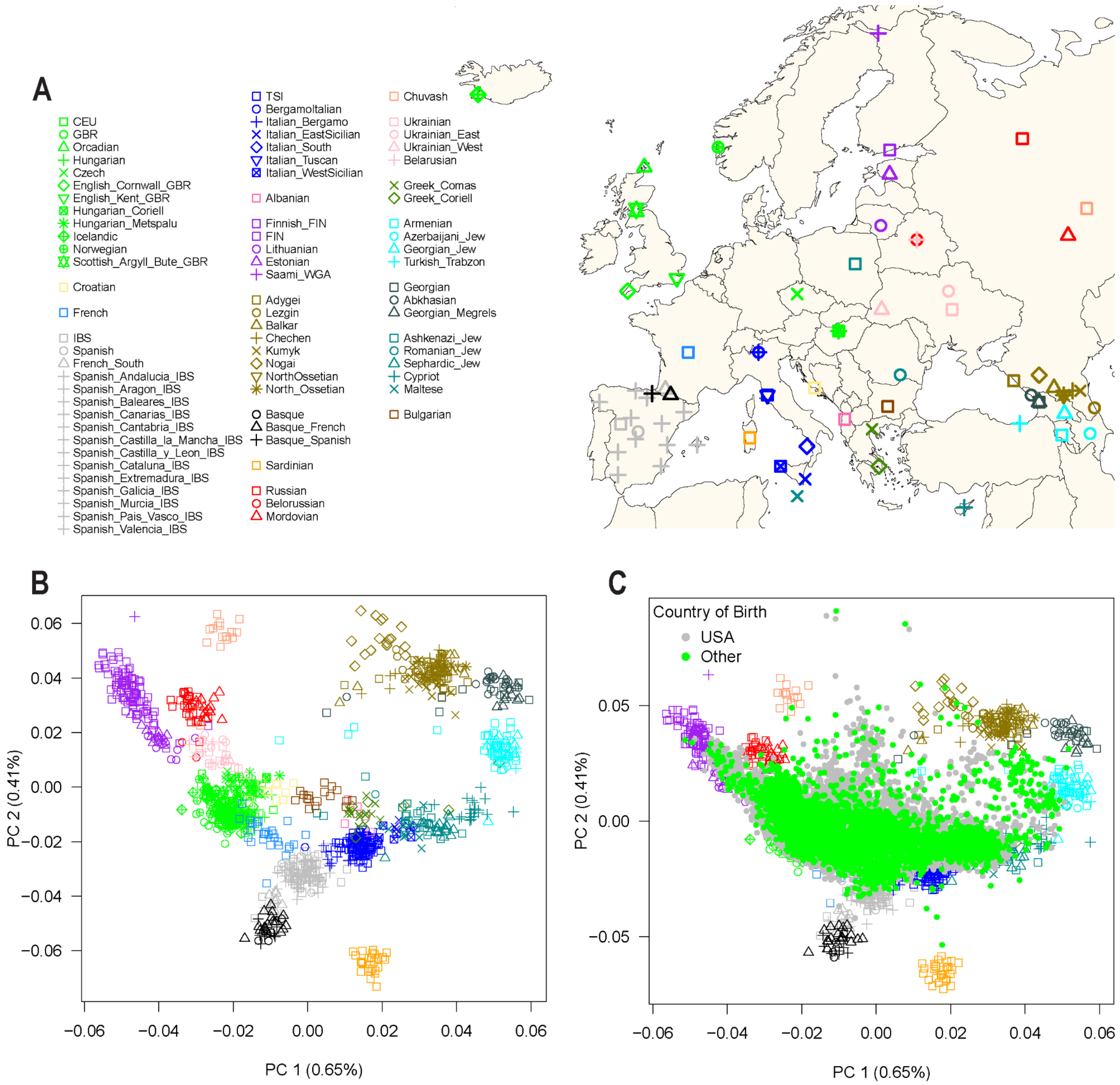


**Fig. S3. European reference panels and coverage of European genetic diversity by *All of Us* participants self-identified as "White".** A) Map of Europe showing the geographic location of samples from 79 European populations. The map was drawn using the R package "maps" version 3.4.1. B) The first two principal components (PCs 1 and 2) of genetic diversity and the percent variance explained. C) Projection of *All of Us* participants self-identifying as "White" onto the European reference panel. This figure is adapted from Gouveia *et al.* 2023 (PMID: 37935687).

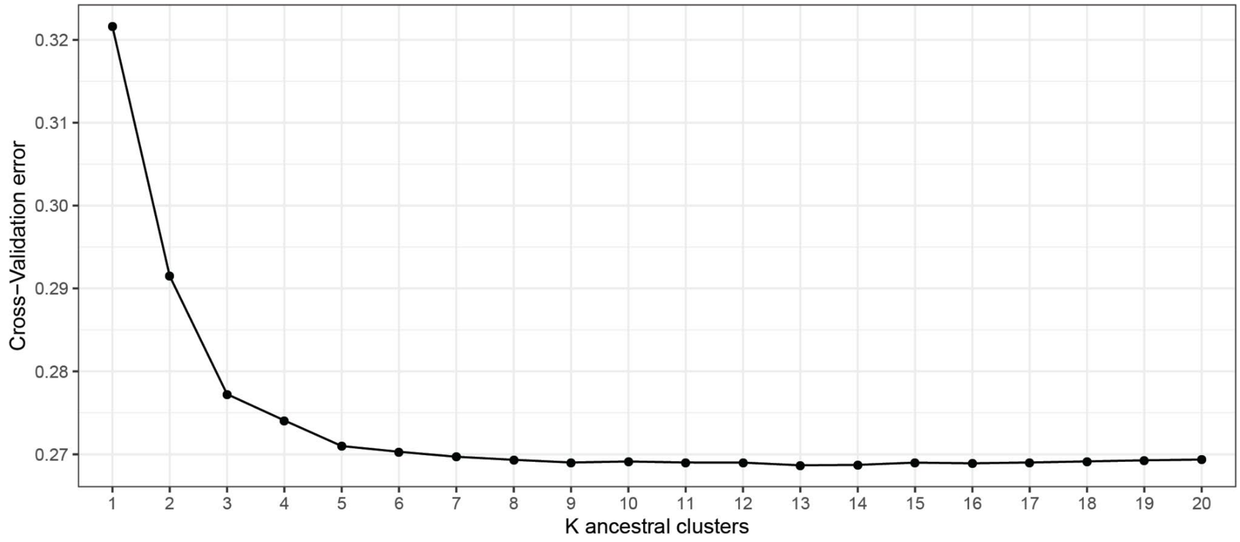


**Fig. S4. Cross-validation plot supporting K = 13 as the most likely number of ancestral components.**
